# A new organotypic model of synaptic competition reveals activity-dependent localization of C1q

**DOI:** 10.1101/118646

**Authors:** Ryuta Koyama, Yuwen Wu, Allison R. Bialas, Andrew Thompson, Christina A. Welsh, Arnaud Frouin, Chinfei Chen, Beth Stevens

## Abstract

Immature neural circuits undergo synaptic refinement, in which activity-dependent competition between synapses results in pruning of inappropriate connections and maintenance of appropriate ones. A longstanding question is how neuronal activity eliminates specific synapses based on their strength. The technical challenges of *in vivo* studies have made it difficult to identify a molecular link between decreased activity and synapse elimination. We developed an organotypic coculture model of the mouse retinogeniculate system that facilitates real-time imaging and elucidation of molecular mechanisms underlying the removal of less active synapses during synaptic competition. Using this model we show for the first time that complement component C1q is necessary for activity-dependent synaptic competition and preferentially localizes to less active, competing presynaptic inputs. In conjunction with classic *in vivo* and *ex vivo* models, this coculture model is a new tool to reveal molecular pathways underlying CNS circuit refinement.

---

As the nervous system matures, neuronal activity sculpts developing synaptic circuits by strengthening certain connections and eliminating others, a process known as synaptic pruning (Kakizawa et al., 2000; Lichtman and Colman, 2000; Mikuni et al., 2013; Penn et al., 1998; Torborg et al., 2005; Yasuda et al., 2011). Pioneering time-lapse imaging of the neuromuscular junction (NMJ) demonstrated that motor neurons engage in a local, activity-dependent competition for postsynaptic territory (Gan et al., 2003; Balice-Gordon and Lichtman, 1994; Buffelli et al., 2003; Lo and Poo, 1991). This competition results in withdrawal of less active inputs and elaboration of the presynaptic termini by the strongest ‘winning’ input (Mooney et al., 1993). Based on these and other studies, it was proposed that activity-dependent molecular cue(s) facilitates specific elimination of less active synapses during competition (Sanes and Lichtman, 1999; Walsh and Lichtman, 2003) however, local competition between CNS synapses *in vivo* has not been visualized or imaged in real-time due to the high density, small size and the difficulty of sparsely labeling individual inputs in the brain. Thus, whether this model is correct or relevant to CNS synapse elimination remains unclear (Hua and Smith, 2004; Huberman, 2007; Katz and Shatz, 1996; Sanes and Lichtman, 1999).

Much of what we know about synaptic pruning in the central nervous system has come from studies using the mouse retinogeniculate synapse as a model system. In this system, axons from retinal ganglion cells (RGCs) project to relay neurons in the dorsal lateral geniculate nucleus (dLGN) of the visual thalamus. Initially, an overabundance of inputs are formed. Over the course of development, these extraneous inputs are eliminated so that a mature relay neuron typically remains innervated by a few strong inputs. Retinogeniculate refinement is dependent on neuronal activity (Penn et al., 1998; Huberman, 2007; Katz and Shatz, 1996; Campbell and Shatz, 1992; Chen and Regehr, 2000; Hong and Chen, 2011; Hooks and Chen, 2006; Mooney et al., 1993; Mooney et al., 1996; Shatz, 1990; Shatz and Kirkwood, 1984; Shatz and Stryker, 1988; Sretavan and Shatz, 1986); however, the molecules that mediate the elimination of specific inputs are poorly understood. If an elimination cue does function during CNS synaptic pruning, we propose that it will localize to synapses undergoing synaptic competition and mediate the elimination of less active synapses.

Complement proteins are a set of secreted innate immune molecules that opsonize, or ‘tag’, cellular debris and pathogens for removal by immune system phagocytes. In addition to this established role in innate immunity, complement is expressed in the healthy brain and is required for developmental synapse elimination (Bialas and Stevens, 2013; Schafer et al., 2012; Stevens et al., 2007). C1q, the initiation protein of the classical complement cascade, is expressed by postnatal retinal ganglion cells (RGCs) and mice deficient in C1q or downstream C3 have sustained defects in synaptic refinement in the retinogeniculate system (Stevens et al., 2007). These data are consistent with a role for C1q as a putative synapse elimination signal; however, whether C1q marks less active synapses undergoing competition with neighboring, more active synapses is unknown.

To investigate these and other mechanistic questions, we developed a novel organotypic coculture model of the retinogeniculate system to image local CNS synaptic competition *in vitro*. In the cocultures, subsets of RGC neurons can be sparsely labeled and neuronal activity in these defined RGC populations can be differentially manipulated, allowing individual inputs competing for postsynaptic territory to be visualized and observed in real-time. Using this model, we provide evidence that local synaptic competition between individual cells mediate CNS synaptic refinement. Moreover, we show that C1q selectively localizes to less active inputs and is required for local activity-dependent synaptic competition. Thus, this coculture system, used in combination with established assays, can elucidate molecular mechanisms of activity-dependent synaptic refinement that have not been possible to resolve *in vivo*.

## Results

### Development of an *in vitro* model of synaptic competition

We developed a coculture model involving retinal explants and dLGN slices to enable visualization and manipulation of individual presynaptic inputs competing for postsynaptic territory. An organotypic slice of the dLGN was prepared from embryonic (E17) mouse embryos and cultured with ventrotemporal retinal explants from E13 embryos electroporated with fluorescent markers and/or other genes of interest (Figure 1a, S1a). The ventrotemporal quadrant of the retina was selectively transfected and cultured because this region has the highest percentage of RGCs that project to the dLGN *in vivo*. (Dhande et al., 2011) We found E17 and E13 embryos promoted optimal survival of relay neurons and RGCs (data not shown). Consistent with *in vivo* studies showing an early developmental window of programmed cell death (Vecino et al., 2004), cleaved caspase-3+ cells were present at DIV 7, rapidly decreased by DIV 14 and was virtually absent by DIV 18 (Figure S1b-d). Furthermore, axon morphology did not show membrane blebbing or fragmentation at or after DIV 14 (Figure S1e). Given that synapse competition peaks shortly after the window of programmed cell death in the retina (Huberman, 2007; Hong and Chen, 2011; Vecino et al., 2004), we focused on DIV 14 cultures or older for subsequent experiments. Cocultures were stained for the relay neuron marker SMI-32 (Bickford et al., 1998; Erişir et al., 1997; Jaubert-Miazza et al., 2005) (Figures 1b-c and S1f, left), the inhibitory neuron marker GAD65 (Figure S1f right), GFAP to stain astrocytes and Iba1 to stain microglia. Healthy RGCs were present in the retina of the cocultures and extend axons into the dLGN (Figure S1a, e). Though some astrocyte processes were observed within the dLGN, most processes were at the edge of the coculture (Figure S1g). Microglia were found throughout the dLGN slice and underwent morphological changes *in vitro* (Figure S1h).

**Figure 1.**
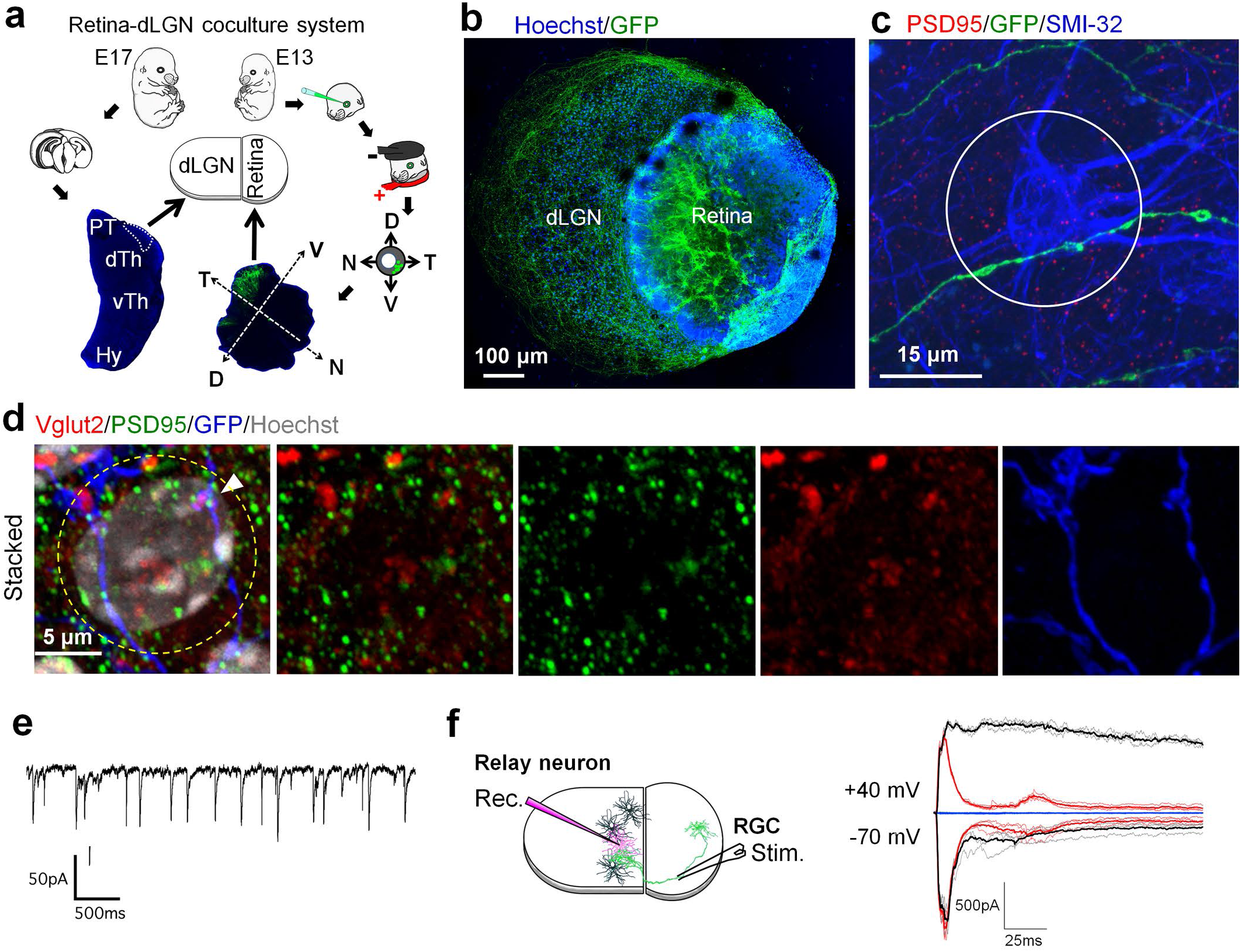
Characterization of a novel retina-dLGN coculture system. (a) Schematic diagram describing the retina-dLGN coculture system. The ventrotemporal region of E13 retinas are electroporated with genes of interest (e.g., GFP or RFP) and cocultured with a dLGN slice microdissected from an E17 mouse embryo. PT, pretectum; dTh, dorsal thalamus; vTh, ventral thalamus; Hy, hypothalamus, V, ventral; N, nasal; D, dorsal; T, temporal. (b) Representative image of a 7 DIV coculture counter-stained with the nuclear marker Hoechst where the retina was transfected with GFP showing axonal growth from the retina into the dLGN slice. (c) Representative image of a GFP+ RGC axon forming *en passant* boutons (swellings) with a SMI-32+ relay neuron. Note the boutons within 15μm of the soma forming *en passant* synapses with the proximal dendrites. (d) Stacked (top) and single plane (bottom) confocal images revealed that axonal swellings colocalize with the RGC presynaptic marker VGlut2 and postsynaptic marker PSD95. (e) Representative spontaneous EPSCs recorded from a relay neuron at -70 mV at 18 DIV. (f) Schematic (left) and representative evoked EPSCs (right, black traces) recorded from a relay neuron in whole-cell patch clamp mode following stimulation of the retinal explant. Synaptic currents were elicited at holding potentials of -70 mV or 40 mV. Addition of the competitive NMDAR antagonist CPP (10μΜ, red traces) led to suppression of the NMDAR-mediated current, indicating the presence of NMDARs at synapses *in vitro*. Further addition of NBQX (10μΜ CPP + 5μΜ NBQX, blue traces) suppressed AMPAR-mediated currents. Stimulus artifacts are blanked for clarity. Darker lines represent the average response.

To identify synapses *in vitro*, cultures were stained with the RGC presynaptic marker VGlut2 and the postsynaptic marker PSD95 (Figure 1d). Axonal swellings were positive for both VGlut2 and PSD95, demonstrating RGCs form *en passant* synapses with relay neurons *in vitro* (Figure 1c, d). EM studies of the intact retinogeniculate system have shown RGCs form synapses along the proximal dendrites of relay neurons (Hamos et al., 1987; Mason, 1982a; Mason 1982b; Raczkowski et al., 1988; Wilson et al., 1984), which we define as dendrites within 15μm of the soma, i.e., dendrites within a circle centered on the soma with radius of 15μm. Few RGCs synapses were detected at distances >15μm from the soma, which is consistent with *in vivo* studies showing most synapses at distal dendrites arise from cortical rather than RGC projections (Erişir et al., 1997) (Figure S1i).

Spontaneous fast excitatory postsynaptic currents (EPSCs) were observed in whole-cell patch clamp configuration of relay neurons from DIV 18 cocultures (Figure 1e), indicating that spontaneous neurotransmission occurs *in vitro*. To investigate whether retinal inputs contribute to functional glutamatergic synapses on relay neurons *in vitro*, synaptic currents were evoked in response to extracellular stimulation of the retinal portion of the coculture, identified by GFP fluorescence. Reliable EPSCs were observed in relay neurons in response to this stimulation (Figure 1f). These currents were inhibited by AMPA- and NMDA-type glutamate receptor antagonists (NBQX and CPP, respectively, Figure 1f). Formation of functional RGC-relay neuron synapses therefore occurs in the cocultures.

### Synapses undergo spontaneous synaptic refinement *in vitro*

Retinogeniculate synapses *in vivo* undergo robust synapse remodeling during development as some inputs strengthen while others weaken and are eliminated. To assess synapse remodeling *in vitro*, we quantified synapse density as the number of PSD95+ axonal swellings of an RGC input divided by the length of the relay neuron dendrite within 15μm of the soma at DIV 14 and 18. Axon length and synapse density averaged across all GFP+ axons were similar at DIV 14 and 18 (Figure 2a, b); however, the distribution of synapses on individual postsynaptic cells shifted over this period (Figure 2c). At DIV 18, there was an increase in low synapse density (<0.05) relay neurons and a decrease in very high synapse density cells (>0.15) compared to DIV 14. These data match the predicted pattern for developmental synaptic remodeling described *in vivo*, that is, a decrease in synapse density from some inputs and an increase in synapse density of the remaining inputs (schematic in Figure 2e). Due to the lack of an optic tract *in vitro*, we cannot simultaneously activate all RCG inputs to quantify the changes in the number of afferent inputs as described in *ex vivo* slices (Hooks and Chen, 2006; Hooks and Chen, 2007). We therefore used a change in the AMPA/NMDA current ratio as a proxy for functional maturation of synapses *in vitro*. We observed a significant increase in the AMPA/NMDA current ratio between DIV 14 and 18 (Figure 2d), consistent with *ex vivo* electrophysiology studies (Hooks and Chen, 2006; Hooks and Chen, 2007). Taken together these data suggest that a form of synaptic refinement occurs *in vitro*. It remains unclear, however, if and how competition between different populations of RGCs contributes to refinement.

**Figure 2.**
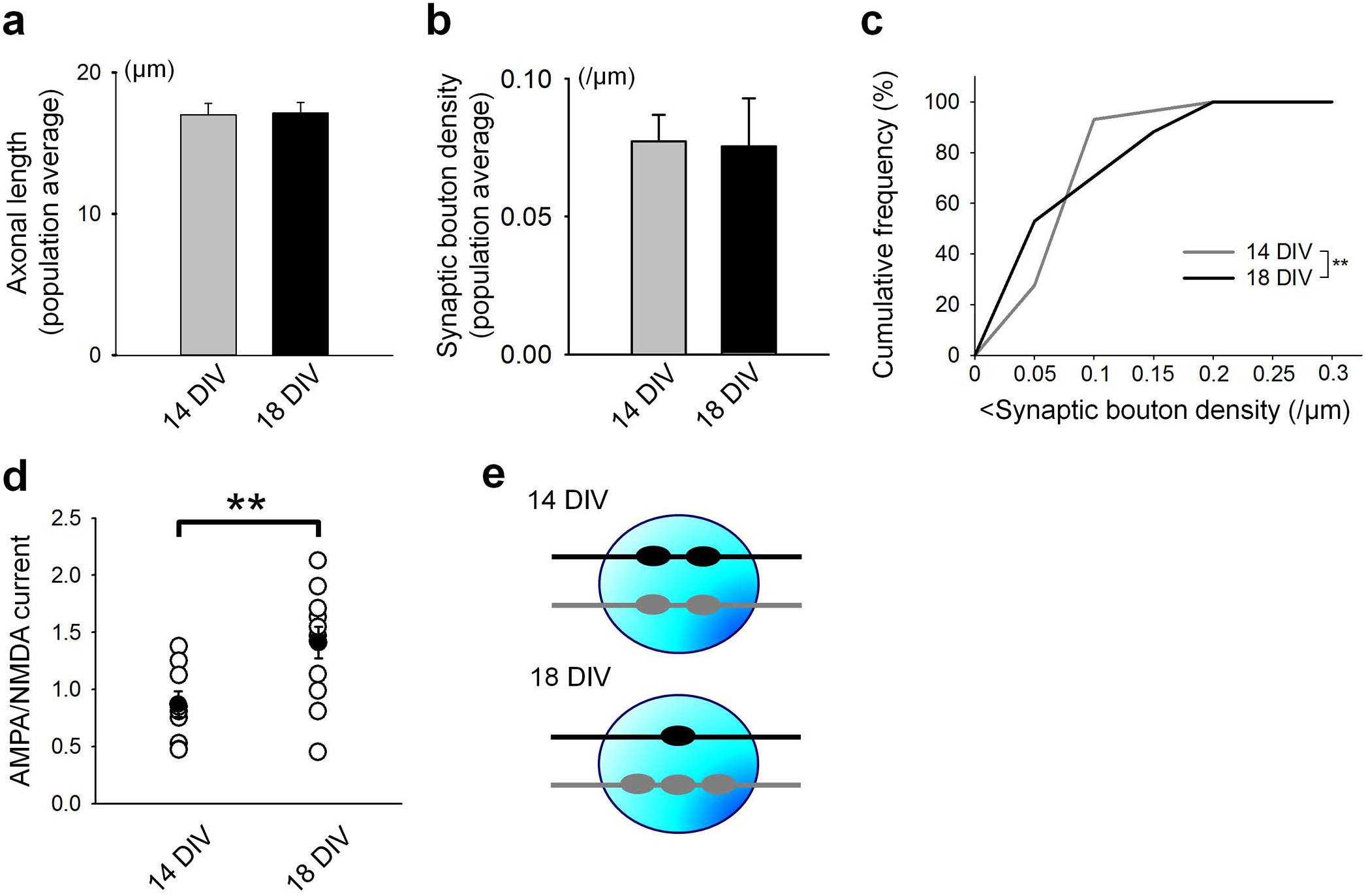
Developmental synaptic remodeling in cocultures. (a) Length of axons within 15μm of the soma did not differ between 14 and 18 DIV. Values were determined by measuring the length of axons within a 15μm radius circle and averaging across all axons regardless of RGC color. 14 DIV: n = 60 axons, 19 cultures. 18 DIV: n = 60 axons, 15 cultures. *P* = 0.911, ns, Student’s t-test. (b) The average density of synaptic boutons across all cells was not significantly different between 14 and 18 DIV. Synapse density was calculated by dividing the number of presynaptic boutons co-localized with PSD95 by the axonal length within a 15μm radius circle around an SMI32+ relay neuron. Values are averages of synaptic density of all RGC axons regardless of color. 14 DIV: n = 20 cells, 10 cultures. 18 DIV: n = 20 cells, 9 cultures. *P* = 0.922, ns, Student’s t-test. (c) Cumulative frequency plots of synapse densities at 14 and 18 DIV. Note the increase in both low and high density inputs at 18 DIV compared to 14 DIV. 4 DIV: n = 29 cells, 11 cultures. 18 DIV: n = 34 cells, 17 cultures. *\*\*P* = 6.71E-05, *F*-test. (d) The AMPA/NMDA current ratio increases between 14 and 18 DIV, suggesting synapse maturation *in vitro*. 14 DIV: n = 9 cells, 5 cultures. 18 DIV: n = 12 cells, 9 cultures. *\*\*P* = 0.00937, Student’s t-test. (e) Schematic of synaptic refinement *in vitro*. RGC axons are represented as black or gray lines, presynaptic boutons as small circles and the relay neuron as the large blue circle.

To determine whether spontaneous synaptic competition between different populations of RGCs occurs *in vitro*, we cultured a dLGN slice with one retina transfected with GFP and a second retina transfected with RFP (Figure 3a, left), allowing us to follow two distinct populations of RGCs as they project to the dLGN (Figure 3a, right). For these experiments, we defined an innervating input as a PSD95+ RGC axonal swelling on a relay neuron and dually innervated cells as individual relay neurons that receive inputs from both a red and a green RGC (Figure 3b). To verify that RGC-relay neuron synapses could be identified using this method, we first stained Hoechst-labeled cocultures with VGlut2 and PSD95. Within this region, axonal swellings in close proximity to single relay neurons were positive for VGlut2 (50/50 boutons, 20 cells) and 92% were also positive for PSD95 (46/50 boutons, 20 cells). To ensure we could observe similar developmental trends, we repeated experiments in figures 2c and 3d in Hoechst-stained cocultures (Figure S3d).

**Figure 3.**
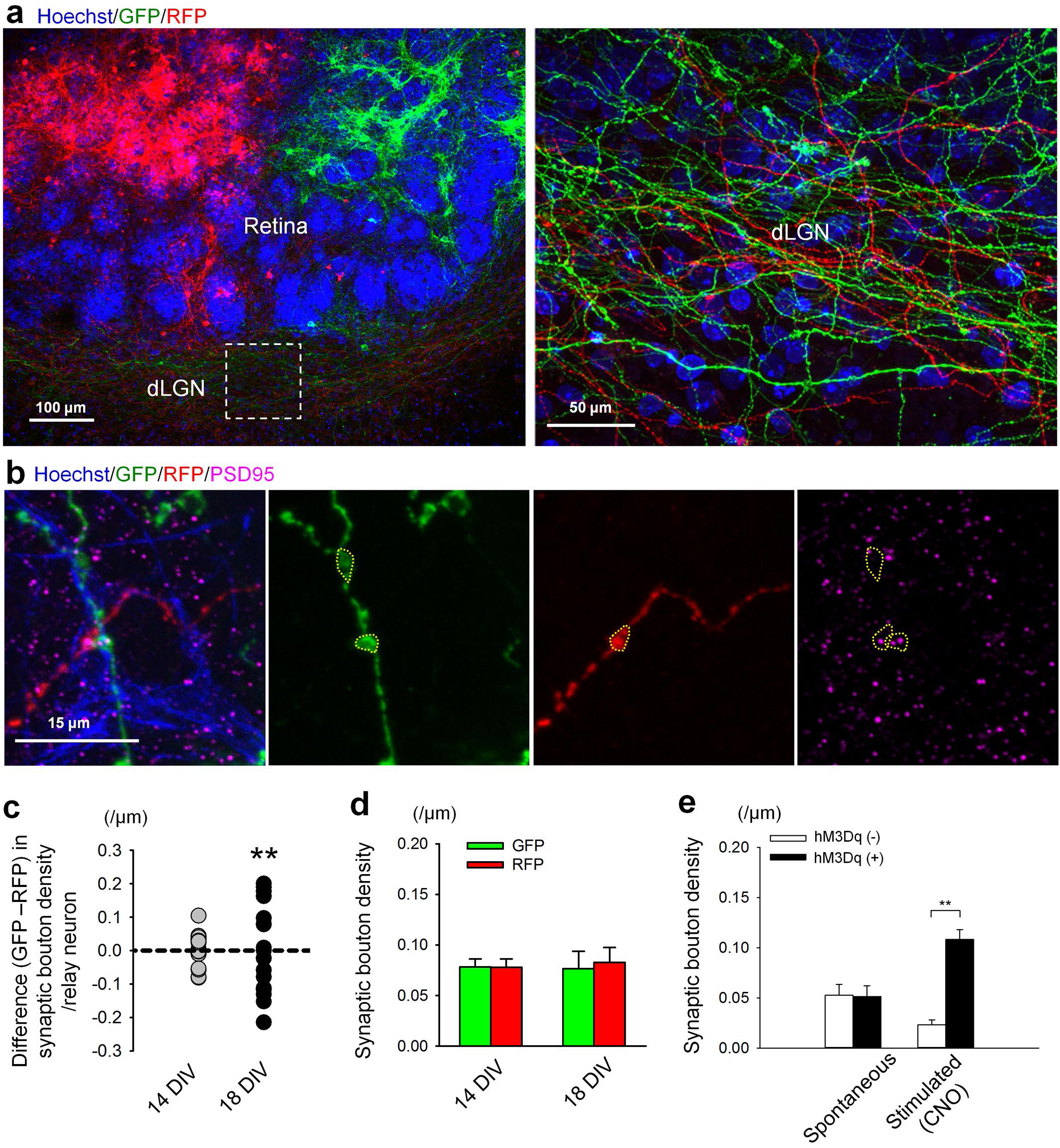
Spontaneous and activity-dependent synapse competition in cocultures. (a) Representative image of a coculture comprised of a GFP-transfected retina and an RFP-transfected retina innervating the dLGN slice (left). The boxed region is magnified on the right and this was the region used for analysis. GFP and RFP axons could be clearly observed running through the dLGN region of the coculture. (b) Representative relay neuron contacted by both a GFP+ and an RFP+ input. (c) Comparison of the number of red and green inputs innervating a single postsynaptic cell at 14 vs.18 DIV under spontaneous conditions. Values close to 0 indicate that both inputs had similar numbers of presynaptic boutons. Most values were clustered close to 0 at 14 DIV, but the distribution of values was significantly wider at 18 DIV. Note the values are equally distributed above and below 0, indicating equal formation of red and green boutons. 14 DIV: n = 20 cells, 9 cultures. 18 DIV: n = 20 cells, 6 cultures. *\*\*P* = 2.81E-05, *F*-test. (d) The total density of synaptic boutons did not differ between the colors (GFP and RFP) at 14 and 18 DIV under basal conditions. 14 DIV: n = 20 cells, 9 cultures. 18 DIV: n = 20 cells, 6 cultures. *P =* 0.988, ns, Tukey-Kramer test following one-way ANOVA. (e) hM3Dq was co-transfected with either RFP+ or GFP+ RGCs. Cultures were treated at 14 DIV for 24 hrs with 10μΜ CNO or vehicle control. Cultures were fixed at 18 DIV and stained for SMI-32 and PSD95. Analysis was performed blinded to treatment condition or hM3Dq transfection status. CNO stimulation led to an increase in hM3Dq+ bouton formation compared to hM3Dq- controls. Spontaneous: n = 21 cells, 12 cultures. Stimulated: n = 26 cells, 20 cultures. One-way ANOVA: *\*\*P* = 6.76E-09. Tukey-Kramer post hoc test: *\*\*P* = 2.90E-09 (hM3Dq- (CNO) vs. hM3Dq+ (CNO)).

We measured synaptic remodeling by quantifying the average density of RGC axon swellings colocalized with PSD95 within 15μm of relay neuron centers at DIV 14 and 18. To then compare RFP+ vs. GFP+ synaptic densities, the RFP+ input density was subtracted from the density of the GFP+ input. Thus, if competition was occurring between these two populations of RGCs (red and green) over this time period, at DIV 14 we would predict similar red and green input densities (GFP-RFP differences close to 0) and at DIV 18 we would predict differing input densities (differences > or < 0).

Consistent with our predictions, at DIV 14, individual relay neurons were frequently dually innervated, with red and green inputs having similar densities on individual relay neurons (Figure 3c). In contrast, at DIV 18, the difference between the density of RFP+ and GFP+ inputs on the same relay neuron was significantly greater than at 14 DIV (Figure 3c). When looking at the entire population of RFP+ and GFP+ axons overall, the average input densities for red and green inputs were unchanged (Figure 3d). These data show that as the cocultures mature, one input gradually increases the number of contacts with a postsynaptic relay neuron, while the other input loses contacts, suggesting local competition between RGC populations contributes to synaptic refinement.

### Relative levels of presynaptic activity bias synaptic competition

As increased bouton numbers have been correlated with increased synaptic strength (Hamos et al., 1987; Hong et al., 2014; Shigemoto et al., 1997), our findings raised the question of whether inputs that are less active are preferentially eliminated while inputs with more activity are maintained during synaptic refinement *in vitro*. To determine whether RGC synaptic competition is dependent on presynaptic activity as described *in vivo*, we chronically stimulated a subset of RGCs using Designer Receptors Exclusively Activated by Designer Drugs (DREADDs) (Armbruster et al., 2007), which are a class of genetically engineered G protein receptors that are activated exclusively by the otherwise biologically inert ligand clozapine-*N*-oxide (CNO). Specifically, we transfected the excitatory form of a modified muscarinic receptor, hM3Dq, into the retinal explant, and assayed the distribution of active (hM3Dq+) versus less active (hM3Dq-) retinal inputs onto individual dLGN relay neurons following CNO stimulation.

hM3Dq has been shown to increase neuronal activity following CNO binding when expressed in certain neurons (Alexander et al., 2009). We verified that treating hM3Dq-transfected retinal explants with CNO increased RGC activity via c-fos immunostaining (Figure S2a, b) and whole-cell RGC recordings (Figure S2c-e). C-fos immunoreactivity significantly increased in GFP+/hM3Dq+ RGCs beginning 1 hour after CNO addition and lasting at least 24 hours, with peak c-fos levels detected at 6 hours (Figure S2a, b). Furthermore, CNO markedly increased the firing rate of hM3Dq+ RGCs (Figure S2c, d), and depolarized the resting membrane potential of these cells (Figure S2e). Together these results confirm that hM3Dq is effective in RGCs.

To determine how driving presynaptic neuronal activity affects synaptic refinement *in vitro*, we transfected retinas with GFP and RFP and co-transfected hM3Dq with either the GFP or RFP in different cohorts. A control and an hM3Dq+ retina were then cultured with a dLGN slice. CNO was applied at DIV 14, a point of robust synaptic remodeling in the cocultures (Figure 2, 3c). After 24 hours, CNO was washed out and replaced with fresh media. Analysis was performed at DIV 18, which represents the end of synaptic competition *in vitro*. Only relay neurons that were dually innervated by both red and green inputs were analyzed.

Cultures treated with CNO showed a shift in input density, with hM3Dq+ inputs showing higher density of PSD95+ axon swellings compared to hM3Dq-controls (Figure 3e: stimulated). In cultures receiving only vehicle, relay neurons were innervated with a comparable number of hM3Dq+ and hM3D1- RGCs at DIV 18 (Figure 3e: spontaneous). Input color alone does not bias synaptic competition, as regardless of the co-transfected fluorophore, the hM3Dq+ RGCs on average outcompeted unstimulated rivals. Furthermore, axon length and branching were not significantly altered in CNO stimulated cultures compared to controls (Figure S2g and S4b), suggesting that axon growth and branching were not affected by CNO at this time point. Our results thus demonstrate that CNO-induced activation of hM3Dq+ inputs increases bouton density of these inputs relative to hM3Dq-inputs.

### Real time imaging of synaptic competition

A major advantage of the coculture system is the ability to visualize the dynamics underlying synaptic competition and remodeling in real time. To investigate this process, we performed time lapse imaging of RGC innervation of relay neurons *in vitro*. As the field lacks tools for live imaging dLGN relay neurons, we found we could use nuclear size as a method to reliably identify relay neurons, as their nuclei are significantly larger than inhibitory neurons in the dLGN (Figure S1f, S3a). Specifically, relay neurons were identified as cells with a nuclear area larger than 73.1μm^2^, which is two standard deviations above the mean area of GAD65+ nuclei (Figure S3a). SMI-32 staining also revealed that relay neuron somas and a portion of the proximal dendrites were reliably found within 7.5μm from the center of the cell (Figure 1c, S3b). We could not use the 15μm radius circle from fixed tissue studies for analysis of time lapse images because, without visualizing the relay neuron processes, it is not possible to determine whether a given input is innervating a given relay neuron. We therefore used a 7.5μm radius circle surrounding the center of Hoechst-stained relay neurons to identify the relay neuron soma and proximal dendrites.

When PSD95+ boutons are analyzed within 7.5μm of either SMI32-stained somas or Hoechst-stained nuclei, we observe a comparable density of synapses regardless of how the relay neuron was identified (Figure S3c). Boutons identified within 7.5μm from a relay neuron nucleus underwent a developmental shift from DIV 14 to 18 (Figure S3e), similar to boutons analyzed in SMI-32 stained cultures. We also observed increased hM3Dq+ bouton density and numbers compared to hM3Dq- bouton density and numbers in CNO treated cultures compared to vehicle treated ones (Figure S3f, g). These experiments confirmed quantification of bouton density using Hoechst as a proxy for the relay neuron is a reliable method to analyze RGC-relay neuron synaptic remodeling.

Cocultures were imaged for up to 24 hours between DIV 15 and 16. RGCs were either transfected with RFP alone or co-transfected with GFP and hM3Dq. Relay neurons were scored as being innervated by a green bouton, a red bouton, boutons of both colors or no boutons at each time point. Cells were grouped into twelve categories based on their innervation status at the beginning and end of imaging (Table S1). Only relay neurons that were innervated by both a red and a green RGC axon at some point during the period of imaging were analyzed to ensure analysis of directly competing inputs. An input was defined as the ‘winner’ of synaptic competition if it maintained a bouton on the relay neuron at the end of imaging and boutons of the rival, ‘loser’ input disappeared before the end of imaging. Although winning and losing RGC inputs could be observed within a short 12-hour imaging window, approximately 30% of cells on average were still dually innervated at the end of imaging under basal conditions, suggesting continued RGC competition for postsynaptic territory at DIV 16 (Table S1).

Consistent with fixed tissue studies (Figure 3), we observed RGC neurons forming mostly *en passant* boutons with relay neurons in the dLGN slice (Figure 4a), although we occasionally observed RGC synapses forming at the axon terminus. In the representative images from a CNO treated culture shown in Figure 4a, a green bouton begins forming at t=0h and has formed by t=5h, while a red bouton formed at hour eight (Figure 4a). This red bouton disappears by hour ten. At the end of imaging (12 hours), only the green bouton is visible, and thus ‘wins’ the competition for this particular relay neuron.

**Figure 4.**
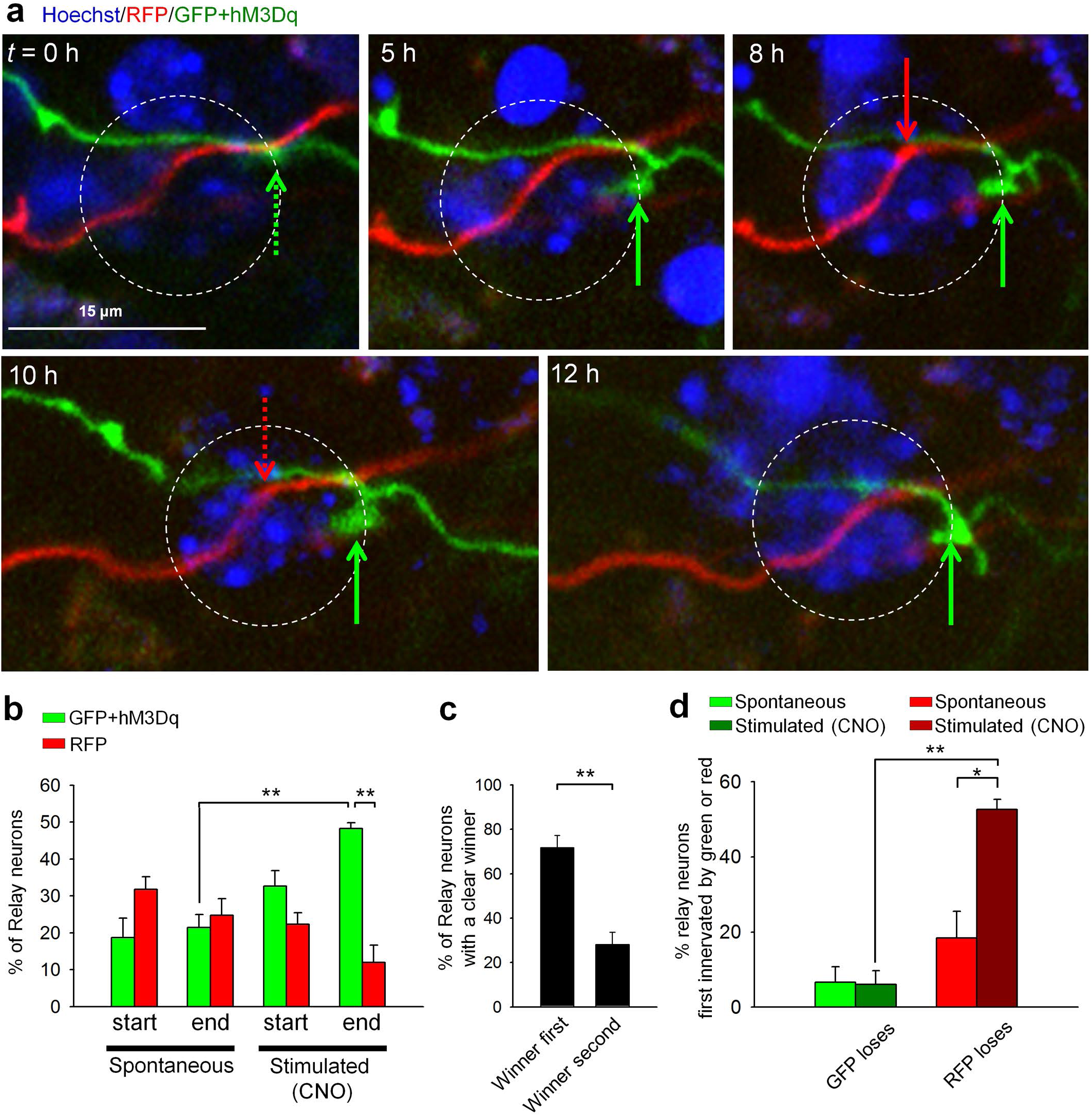
Time lapse imaging of activity-dependent synaptic competition in cocultures. (a) Representative time lapse images of synaptic competition *in vitro* following 12hrs of CNO stimulation. Arrows denote presynaptic boutons, and dashed arrows indicate appearing or disappearing boutons. CNO (10μΜ) was added 12hrs before imaging as well as directly into the imaging media for a total of 24hrs of stimulation. The culture shown here was imaged once every hour for 12hrs. (b) Quantification of % of relay neurons per culture analyzed which were innervated by either a GFP+/hM3Dq+ or RFP+ bouton at the beginning of imaging and at the end of imaging under spontaneous (-CNO) or stimulated (+CNO) conditions. % relay neurons per culture were averaged for final estimates. There was no difference between the proportion of relay neurons innervated by green or red boutons at the beginning of imaging with or without CNO, or at the end of imaging without CNO. Following CNO addition, there was a significant increase in the proportion of relay neurons innervated by GFP+/hM3Dq+ boutons compared to RFP+ boutons at the end of imaging. There was also a significant increase in green boutons at the end of imaging following CNO stimulation compared to number of boutons at the end of imaging under spontaneous conditions. Analysis was performed on all dually innervated cells. Spontaneous: n = 109 cells, 5 cultures. Stimulated: n = 104 cells, 4 cultures. One-way ANOVA: *\*\*P* = 6.84E-05. Tukey-Kramer post hoc test: *\*\*P* = 0.00135 (GFP/hM3dq (spontaneous, end) vs. GFP/hM3Dq (CNO, end)), *\*\*P* = 3.79E-05 (GFP/hM3Dq (CNO, end) vs. RFP (CNO, end)). (c) Under spontaneous conditions, inputs that first formed a presynaptic bouton on a relay neuron were significantly more likely to persist and be maintained through the imaging session compared to inputs that formed a bouton second. Analysis was performed on cells first innervated by either a red or green bouton and where a clear winner emerged at the end of imaging. Winner first (winning inputs arrived first): n = 22 cells, 5 cultures. Winner second (winning inputs arrived second): n = 8 cells, 5 cultures. *\*\*P* = 4.75E-04, Student’s t-test. (d) GFP+/hM3Dq+ inputs that arrived first rarely lost competition regardless of treatment condition, while RFP+ inputs that arrived first in non-stimulated cultures also rarely lost competition. However, RFP+ inputs significantly lost competition to CNO-activated GFP+/hM3Dq+ inputs, even when the latter arrived after the RFP+ input. Analysis was performed on relay neurons that were first innervated by a green bouton (GFP loses) or first innervated by a red bouton (RFP loses). GFP loses, Spontaneous: n = 2/20 cells, 5 cultures. GFP loses, Stimulated: n = 2/33 cells, 4 cultures. RFP loses, Spontaneous: n = 6/35 cells, 5 cultures. RFP loses, Stimulated: n = 13/24 cells, 4 cultures. One-way ANOVA: *\*\*P* = 0.001207. Tukey-Kramer post hoc test: *\*\*P* = 0.00169 (GFP loses (CNO) vs. RFP loses (CNO)), *P = 0.0164 (RFP loses (Spontaneous) vs. RFP loses (CNO)).

To investigate the role of presynaptic activity in synaptic competition in real time, cultures were treated for 24 hours with 10μΜ CNO (‘stimulated’ samples) as was done for fixed tissue studies. Control cultures were untreated (‘spontaneous’). In both spontaneous and stimulated cultures, relay neurons were innervated by approximately equal numbers of RFP+ and GFP+/hM3Dq+ inputs at the beginning of imaging (Figure 4b). At the end of imaging, relay neurons in spontaneous cultures were innervated by similar levels of RFP+ and GFP+/hM3Dq+ inputs, but an increase in GFP+/hM3Dq+ boutons vs. RFP+ boutons was observed in cultures stimulated by CNO (Figure 4b). Furthermore, there is an increase in GFP+/hM3Dq+ bouton density after CNO stimulation compared to unstimulated controls at the end of imaging (Figure 4b). These data are consistent with our fixed tissue studies. There was also a significant increase in number of relay neurons where the GFP+/hM3Dq+ input ‘won’ at the end of imaging following CNO stimulation compared to number of GFP+/hM3Dq+ winners under spontaneous conditions.

In stimulated cultures we observed both increased formation of GFP+/hM3Dq+ boutons (~30% green boutons at t = 0 h to ~50% green boutons at the end of imaging) as well as increased disappearance of RFP+ boutons (~20% red boutons at t = 0 h to ~10% red boutons at the end of imaging) compared to unstimulated controls. Increased synaptogenesis of green inputs and increased elimination of red inputs likely both contribute to CNO-induced skewing of synaptic competition. These results are consistent with other studies showing synapse formation and elimination occur simultaneously during circuit refinement (Hua and Smith, 2004; Sanes and Lichtman, 1999; Balice-Gordon and Lichtman, 1994; Armbruster et al., 2007; Alexander et al., 2009; Cline, 2001; Goda and Davis, 2003; Trachtenberg et al., 2002).

### Timing of bouton formation biases synaptic competition

Studies at the NMJ and in the cerebellum suggest that synapses that form first are generally maintained over those that arrive later (Gan et al., 2003; Balice-Gordon and Lichtman, 1994; Buffelli et al., 2003; Turney and Lichtman, 2012). We asked whether this rule also applies to RGC bouton formation, a question that could not previously be addressed without live imaging of RGC synaptic competition. Of the 30 total relay neurons (5 cultures) where a single RGC ‘arrived’ first and where a clear ‘winner’ emerged at the end of imaging, 22 were innervated by the same input at the end of imaging, while only 8 of these cells were innervated by an input that arrived second (Figure 4c, Table S1). Inputs that form a synapse before a rival cell are thus at a clear advantage during spontaneous synaptic competition.

To determine the relative contribution of input timing and neuronal activity in biasing the outcome of synaptic competition, we analyzed relay neurons that were initially innervated by a single input of either color in stimulated cultures compared to spontaneous control cultures. Interestingly, in stimulated cultures, if a CNO-activated GFP+/hM3Dq+ input arrived after a RFP+ input, the RFP+ bouton significantly lost competition for the relay neuron (13 out of 24 cells, Figure 4d). In contrast, of the inputs first innervated by RFP+ axons, only 6 out of 35 lost competition under spontaneous conditions (Figure 4e, Table S1: column 6). When GFP+/hM3Dq+ inputs were the first to arrive, RFP+ inputs were able to outcompete the green axons in only 2 out of 20 cells (spontaneous) and 2 out of 33 cells (stimulated). Thus, GFP+/hM3Dq+ inputs arriving first rarely lost competition regardless of whether they were or were not treated with CNO (Figure 4d, Table S1: column 8). These data suggest that during synaptic competition there is a preference for inputs that arrive first, although axons with increased levels of activity relative to their neighbors gain a significant competitive advantage.

### C1q is required for activity-dependent synaptic competition

Previous work demonstrated that C1q is required for developmental pruning in the retinogeniculate system in vivo (Bialas and Stevens, 2013; Stevens et al., 2007). However, these studies did not distinguish whether C1q regulates activity-dependent elimination of locally competing inputs. To address this question, E13 C1q knockout (KO) retinas were transfected with either GFP/hM3Dq or RFP and cultured with a dLGN slice from an E17 C1q KO; control cultures were made from wild type (WT) tissue. Consistent with the role of C1q in developmental pruning, C1q KO cocultures had significantly higher synapse densities overall compared to WT cocultures at 18 DIV (Figure S4a). Following CNO application, there was no significant difference between GFP+/hM3Dq+ synapse density and RFP+ synapse density in C1q KO cocultures, in contrast to WT cocultures (Figure S4a). However, the increased numbers of synapses in KO cultures under spontaneous conditions could occlude an activity-dependent shift.

To gain temporal control and manipulate C1q solely during the period of synaptic competition *in vitro*, we used a specific C1q function blocking antibody, ANX-M1 (Annexon Biosciences), which has been shown to neutralize C1q activation of the classical complement cascade (Hong et al., 2016). ANX-M1 or an isotype control was added to the cocultures at DIV 14. The hM3Dq was randomly co-transfected in either RFP+ or GFP+ retinas, and CNO or vehicle control was applied at DIV 14. CNO or vehicle was washed out after 24 hours with fresh media and ANX-M1 or the isotype control was re-applied until DIV 18. Importantly, no differences in hM3Dq+ vs. hM3Dq-densities were observed in ANX-M1 treated cocultures vs. isotype treated cultures under spontaneous conditions when vehicle control was applied (Figure 5a). In contrast, under stimulated CNO-treated conditions, a significant increase in hM3Dq+ densities was observed in isotype treated cultures, but not in ANX-M1 treated cultures (Figure 5a). We did not observe differences in axon branching between the different treatment condition (Figure S4b), indicating that C1q is necessary for local activity-dependent synaptic competition.

**Figure 5.**
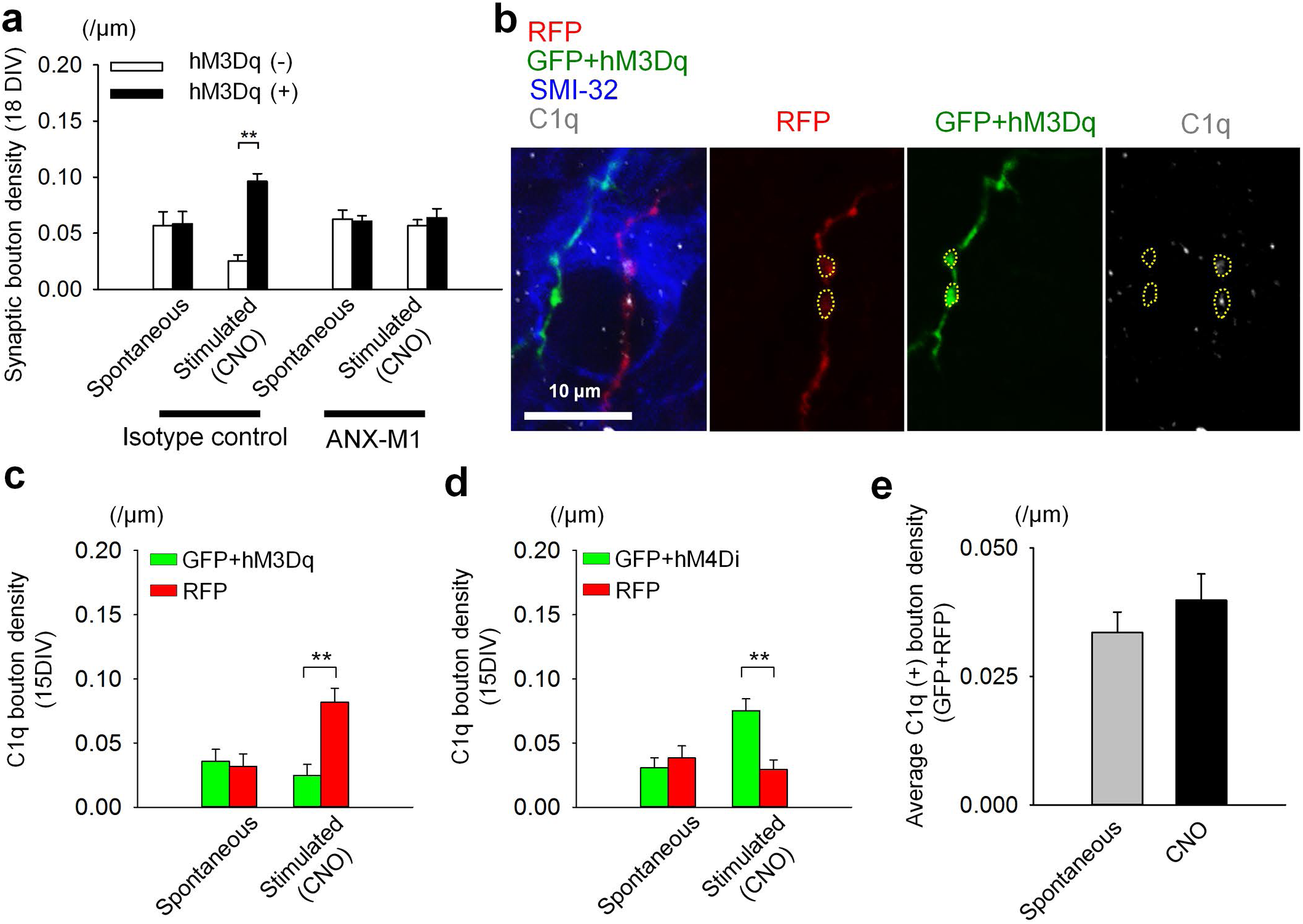
C1q is necessary for synaptic competition and preferentially labels less active inputs. (a) WT cocultures were treated with or without CNO at 14 DIV for 24hrs. Cultures were also treated with 15μg/ml of ANX-M1 or an isotype control at 14 DIV. Following wash-out of CNO, ANX-M1 or the isotype control was re-applied until 18 DIV, when the cultures were fixed and stained for PSD95 and SMI-32. hM3Dq was transfected in either the red or green RGCs and the experimenter was blinded to treatment condition and hM3Dq transfection during analysis. CNO stimulation led to an increase in hM3Dq+ boutons in isotype treated cultures, but no increase was seen in ANX-M1 treated cultures. No increase in baseline bouton density was observed in either treatment condition. Isotype, - CNO: n = 25 cells, 15 cultures. Isotype, +CNO: n = 28 cells, 24 cultures. ANX-M1, -CNO: n = 28 cells, 24 cultures. ANX-M1, +CNO: n = 25 cells, 20 cultures. One-way ANOVA: *\*\*P* = 1.92E-06. Tukey-Kramer post hoc test: *\*\*P* = 2.04E-08 (hM3Dq- (Isotype control, CNO) vs. hM3Dq+ (Isotype control, CNO)). (b) Representative confocal images of a relay neuron innervated by both GFP+/hM3Dq+ and RFP+ axons at 15 DIV. CNO (10μΜ) was applied at 14 DIV for 18hrs. (c, d) C1q preferentially co-localizes with less active inputs after CNO stimulation. In GFP+/hM3Dq+ and RFP+ cultures, RFP+ inputs had increased labeling. The reverse was true in GFP+/hM4Di+ and RFP+ cultures. Spontaneous (hM3Dq): n = 12 cells, 4 cultures. Stimulated (hM3Dq): n = 19 cells, 6 cultures. One-way ANOVA: *\*\*P* = 8.15E-05. Tukey-Kramer post hoc test: *\*\*P* = 0.000200. Spontaneous (hM4Di): n = 22 cells, 7 cultures. Stimulated (hM4Di): n = 27 cells, 0 cultures. One-way ANOVA: **P = 0.000346. Tukey-Kramer post hoc test: *\*\*P* = 0.000346. (e) Average C1q+ bouton density of all inputs (both red and green) in non-stimulated and stimulated cultures were not significantly different. Spontaneous: n = 28 cells, 12 cultures. Stimulated: n = 46 cells, 26 cultures. *P =* 0.393, ns, Student’s t-test.

### C1q preferentially labels competing less active synapses

What is the mechanism of C1q-mediated synaptic competition? C1q could selectively localize to less active synapses and lead to specific elimination of those synapses by microglia or other mechanisms. Alternatively, C1q could target synapses more broadly, and a protective mechanism could maintain or protect more active synapses from elimination. To determine whether C1q preferentially localizes to less active inputs, DIV 14 WT cocultures were treated with CNO or vehicle for 18 hours and then immunostained for C1q (Stephan et al., 2013). Dually innervated relay neurons were analyzed for density of C1q+ boutons (Figure 5b-d). GFP+ inputs were co-transfected with either hM3Dq to activate the green RGCs or with hM4Di to inhibit activity. When only vehicle control was applied, C1q was detected at RFP+ and GFP+ boutons at low levels and with equal frequency at DIV 15 (Figure 5c, d: spontaneous). However, after CNO application, less active inputs (hM3Dq- inputs (Figure 5c: stimulated) and hM4Di+ inputs (Figure 5d: stimulated)) were consistently labeled with more C1q relative to their more active rivals. Similar results were observed using Hoechst to label relay neurons (Figure S4d, e).

The pattern of C1q immunostaining is similar to what has been observed *in vivo* in the developing dLGN, with C1q puncta localized to cell bodies, axons, dendrites and synaptic boutons and about 20% of VGlut2 puncta co-localized with C1q puncta (Bialas and Stevens, 2013). No C1q staining was observed in KO cocultures (Figure S4c), indicating the specificity of the antibody. The overall C1q+ bouton density in vehicle and CNO-treated cocultures was not significantly different (Figure 5e), indicating that C1q distribution seems to change such that it co-localizes more with hM3Dq- boutons compared to hM3Dq+ boutons. Taken together, these results suggest that C1q is required for local synaptic competition and reveal for the first time that C1q can selectively localize to less active synapses.

## Discussion

In this study, we developed a new organotypic coculture system to investigate molecular mechanisms underlying CNS synaptic competition and remodeling that have been difficult to address *in vivo*. Time-lapse imaging of CNS synapses can be performed in the cocultures, revealing that local competition between presynaptic inputs mediates CNS synaptic refinement. Furthermore, both timing of input innervation and presynaptic activity influence the outcome of synaptic competition. We also determined that C1q is necessary for the local activity-dependent synaptic competition we observed via time lapse imaging. Finally, we show for the first time that C1q preferentially localizes to less active presynaptic inputs undergoing competition, consistent with a role as a synapse elimination cue. This *in vitro* model therefore enables elucidation of molecular mechanisms underlying CNS circuit refinement that, when used in parallel with established *in vivo* models, may be broadly relevant to other sensory systems and brain regions.

### An *in vitro* model of synaptic competition

Due to the difficulty of investigating mechanisms of synaptic competition and refinement *in vivo*, several *in vitro* models have been developed; however, our model has some unique advantages. Early work in the retinogeniculate system using *ex vivo* acute slice cultures revealed the contribution of spontaneous retinal activity to retinogeniculate synapse development (Penn et al., 1998; Katz and Shatz, 1996; Campbell and Shatz, 1992; Mooney et al., 1993; Mooney et al., 1996; Shatz, 1990; Shatz and Kirkwood, 1984; Shatz and Stryker, 1988; Sretavan and Shatz, 1986; Stellwagen and Shatz, 2002; Stellwagen et al., 1999). Although these models contributed greatly to our understanding of activity-dependent refinement, visualization of synaptic competition and manipulation of distinct presynaptic populations is difficult in acute slices. An organotypic model of the kitten retinogeniculate system has been described (Guido et al., 1997) and provided rationale for a rodent model that will enable genetic and transgenic approaches to manipulate C1q and other molecular candidates. Previous mouse coculture systems of the visual pathway focused on neurite outgrowth rather than synaptic refinement and competition (Smalheiser, 1981; Washburn et al., 2011). Organotypic coculture models of the cerebellum (Letellier et al., 2009; Uesaka et al., 2012; Mikuni et al., 2013) and corticospinal tract (Ohno et al., 2004; Ohno et al., 2010) have been developed to observe synaptic remodeling in other CNS regions. However, in these systems, only a single population of inputs was analyzed.

To determine if C1q targets less active (loser) inputs competing against more active (winner) inputs, we needed a system that allows identification and manipulation of two distinct populations of presynaptic cells. We achieved such a system by culturing two separate retinal explants with a single dLGN slice. Spontaneous synaptic remodeling occurs in the cocultures, and when one population of RGCs was specifically activated relative to the other population, competition was skewed in favor of the activated inputs. We observed this phenomenon via both fixed tissue studies and live imaging of RGCs directly competing to innervate relay neurons. This competition and the mechanisms we describe could describe inter-eye or intra-eye synaptic competition, and further investigation both *in vitro* and *in vivo* will help to distinguish between these two processes. The coculture system is thus a unique and valuable *in vitro* model of CNS synaptic competition that can be used in combination with *in vivo* models.

### C1q preferentially targets less active synapses during synaptic competition

Previous *in vivo* studies in the retinogeniculate system identified C1q as a putative molecular elimination signal (Bialas and Stevens, 2013; Schafer et al., 2012; Stevens et al., 2007). However, whether C1q mediates local activity-dependent synaptic competition via ‘tagging’ of less active inputs was unknown due to the difficulty of imaging single competing CNS synapses *in vivo*. Here, we first showed RGCs engage in local competition for relay neuron ‘territory’ in an activity-dependent manner, with more active inputs outcompeting less active inputs. When C1q was then eliminated from cocultures either via culturing from KO mouse tissue or via pharmacological inhibition, more active inputs no longer outcompeted less active inputs, demonstrating for the first time that C1q is required for RGCs to engage in activity-dependent synaptic competition. Furthermore, as the cocultures allow visualization of dually innervated relay neurons, we demonstrated that C1q preferentially colocalizes with less active inputs undergoing direct competition with more active synapses.

Combined with studies of C1q *in vivo*, our findings thus suggest that C1q mediates developmental pruning by localizing to less active synapses. Interestingly, as C1q KO cocultures showed increased baseline synapse density when compared to WT cultures, C1q could also be important for synaptogenesis or axonal pruning. This is further supported by the many non-synaptic C1q puncta observed in the cocultures, which is consistent with what has been previously described in the retinogeniculate system *in vivo* (Bialas and Stevens, 2013). The role of CNS C1q outside of synaptic competition and pruning is still unexplored, and the cocultures could aid in elucidating these and other pathways.

Another important open question from these studies is how C1q is able to preferentially localize to competing, less active inputs. One possibility is that C1q is specifically recruited to less active synapses by binding a factor or receptor enriched at less active synapses. The coculture model we describe here will aid identification of putative complement binding proteins due to the ability to visualize and label individual competing inputs. Alternatively, C1q could be excluded from more active synapses by protective molecules, as proposed in the punishment model (Sanes and Lichtman, 1999; Lichtman and Colman, 2000). These possibilities are not mutually exclusive and will be important questions for future investigations. Finally, both microglia and astrocytes have been shown to be involved in developmental pruning *in vivo* (Schafer et al., 2012; Chung et al., 2013), and the coculture system will allow development and optimization of novel *in vitro* assays to interrogate these mechanisms in future studies.

Understanding the developmental mechanisms of synapse remodeling has broad implications for understanding CNS pathology, as these mechanisms could also mediate abnormal circuit refinement in neuropsychiatric disorders or aberrant loss of synapses in neurodegenerative diseases. Complement proteins in particular have recently been implicated in pathological synapse elimination in schizophrenia (Sekar et al., 2016), Alzheimer’s disease (Hong et al., 2016), glaucoma (Williams et al., 2016), frontotemporal lobar degeneration (Lui et al., 2016) and other diseases that involve synapse loss and dysfunction. Moreover, other molecular pathways such as MHC class I molecules (Corriveau et al., 1998; Goddard et al., 2007; Huh et al., 2000; Lee et al., 2014) are known to mediate proper circuit refinement and contribute to synapse loss in aged and disease brains. In conjunction with *in* vivo models, these and other signaling pathways could be investigated in the cocultures to reveal novel interactions or complementary roles underlying synapse refinement. The coculture model described here would therefore provide an invaluable addition to the toolbox for investigating CNS synapse refinement and elimination in development and disease.

## Materials & Methods

### Animals

C57BL/6 mice were obtained from Charles River. *C1qa-/-* mice (C57BL6 background) were generously provided by M. Botto (Botto et al., 1998). All experiments were approved by the Boston Children’s Hospital institutional animal use and care committees and performed in accordance with all NIH guidelines for the humane treatment of animals.

### Pharmacological agents

Clozapine-N-oxide (CNO) was purchased from Sigma-Aldrich. ANX-M1 and its isotype control were provided by Annexon Biosciences (Hong et al., 2016).

### Preparation of plasmid DNA-micropipettes

Fine-tipped glass micropipettes (PCR pipette, Drummond, 5-000-1001-X10) were pulled and the tip was gently broken to make an angled point. DNA solutions were prepared to desired concentrations as follows with the addition of Fast Green Dye (0.05%) to visualize injections: membrane-targeted GFP or tdTomato from Clontech, 6 μl DNA solution (1-3 μg/μl) in TE buffer + 0.5 μl Fast Green Dye; for co-transfection of membrane-targeted GFP and hM3Dq (generous gift from Dr. Bernardo Sabatini), 4 μl of hM3Dq plasmids (~2 μg/μl) + 2 μl of membrane-targeted GFP (~2 μg/μl) + 0.5 μl Fast Green Dye. DNA solution was aspirated into the pipette through an aspirator tube (15’’ Aspirator Tube Assembly, 2-000-000). The DNA-containing micropipettes were kept at room temperature before use.

### Ex vivo retinal electroporation and the preparation of retinal explants

E13.5-E14.5 pregnant mice were anesthetized with isoflurane, embryos were decapitated and their heads collected in ice-cold Hanks’ balanced salt solution (HBSS). Plasmid DNA (0.3-0.5μl) was injected into retinas via aspiration of DNA solution from glass pipettes such that the tip of the pipette entered the retina from the nasal region of the eye. Electroporation (Tweeztrods, 7mm, Harvard apparatus, 450165) was performed such that the positive electrode was adjacent to the ventral side of the head. Currents were delivered using an ECM 830 electroporator with the following settings: Mode, LV; Voltage, 45 V; Pulse length, 50 mS; # of pulses, 4; interval, 100 ms. After electroporation, retinas were microdissected from the head and cultured in serum-free RGC culture medium (Bialas and Stevens, 2013; Meyer-Franke et al., 1995) without growth factors for 24–48hrs at 37°C, 10% CO_2_. Subsequently, the fluorescent regions of the retina were dissected out in Dulbecco’s phosphate-buffered saline (Invitrogen 14287-080) under a fluorescent dissection microscope equipped with P-FLA2 fluorescence attachment 2 (Nikon). Non-fluorescent portions of the retina were discarded.

### Cocultures of retinal explants and dLGN slices

E17 mouse brains were sectioned into 350-μm-thick coronal slices using a Mc ILWAIN tissue chopper, and the slices that include diencephalic regions were collected in ice-cold HBSS. Caudal sections including the optic tract, pretectum (PT) and dorsal and ventral thalamus (dTh, vTH) were selected (see Figure 1A, S1B). In these caudal thalamic slices, the thalamic radiation, which forms the medial border of the lateral geniculate, was visible, and the dLGN region was microdissected out. The dLGN slices were incubated for 30-90 min at 4°C in incubation medium containing minimal essential medium (MEM), 10 mM Tris, 25 mM HEPES, and 60 mM glucose supplied with penicillin and streptomycin. Following this incubation, retina(s) were attached to the lateral side of a dLGN slice on a membrane filter (Millicell-CM PICMORG50; Millipore), which was coated with poly-D-lysine (Sigma, final: 10 μg/ml), laminin (R&D, final: 2 μg/ml), and rat tail collagen solution (Gibco, final: 50 μg/μl). The cocultures were cultured with serum-free RGC culture medium (Bialas and Stevens, 2013; Meyer-Franke et al., 1995) supplied with 76 mM glucose at 37°C, 10% CO_2_. 5% FBS was included in the media for only the first 3 DIV, and the medium was changed every 3 days with serum-free media. To stimulate hM3Dq+ neurons, 10μM CNO was applied to the culture media (see text for duration of CNO stimulation for each experiment). For the C1q blocking experiments, ANX-M1 or isotype control were added at 15μg/ml as previously described for acute slices (Hong et al., 2016).

### Immunostaining and confocal imaging

Cultured samples were fixed in 4% PFA at 4°C for 24hrs. For C1q staining, the fixation was performed at room temperature for 30 min. The fixed samples were rinsed 3 times with 0.1 M phosphate buffer (PB). Slices were then permeabilized and blocked for 1hr at room temperature in 0.1 M PB + 0.3% Triton X-100 with 10% goat serum. The samples were subsequently incubated with primary antibodies in 0.1 M PB + 0.3% Triton X-100 with 10% goat serum at 4°C for 48hrs with agitation. Cultures were rinsed 3 times with 0.1 M PB and then incubated with secondary antibodies in 0.1 M PB + 0.05% Hoechst + 0.3% Triton X-100 with 10% goat serum at 4°C overnight with agitation. After the rinse, samples were embedded in Fluorescent Mounting Medium (Dako, S3023).

The following primary antibodies were used for immunostaining: chicken anti-GFP (1:500; Abcam, ab13970), rabbit anti-c-fos (1:500; Santa Cruz, sc52), rabbit anti-PSD95 (1:500; Zymed/Invitrogen, 51-6900), mouse anti-SMI32 (1:500; Covance, SMI-32R), guinea pig anti-Vglut2 (1:500; Millipore, AB2251), mouse anti-MAP2 (1:500; Sigma, M1406), mouse anti-NeuN (1:500; Millipore, MAB377), rabbit anti-GAD67 (1:500; ab97739) rabbit anti-cleaved caspase-3 (1:500; cell signalling, 9661S), rat anti-RFP (1:500; Allele Biotechnology, ACT-CM-MRRFP10), and rabbit anti-C1qA (Epitomics, custom antibody synthesis, undiluted culture supernatant, validated in (Stephan et al., 2013). The following secondary antibodies were used: Alexa Fluor 488-, 594-, or 647-conjugated secondary antibodies (1:500; Invitrogen).

For analysis of synapse density, images were acquired using an LSM700 scanning confocal microscope equipped with diode lasers (405, 488, 555 and 639nm) and Zen 2009 image acquisition software (Carl Zeiss). Z-series images were collected with a 1.4NA 63× oil immersion objective at a voxel size of 0.0990.099-0.2 μm (x-y-z). C1q immunostained images and SMI-32-stained images were acquired using an UltraView Vox spinning disk confocal microscope equipped with diode lasers (405nm, 445nm, 488nm, 514nm, 561nm, and 640nm) and Volocity image acquisition software (Perkin Elmer). Z-stacks were acquired such that the entire thickness of the LGN innervated by RGC axons was captured in the series. All analyses were performed blinded to experimental condition.

### Quantification of synapse density

Cells with a nuclear area greater than 73.1μm^2^ were identified as relay neurons. A presynaptic input was defined as a single RGC axon, while axonal swellings were identified by eye and defined as presynaptic boutons. These swellings were distinct from degenerating axons as the latter were characterized by loss of the axon and small, bright puncta corresponding to axonal debris (see Figure 1d, middle panel). Boutons that colocalized with the postsynaptic markers PSD95 or Homer were defined as synapses and co-localization was confirmed by threedimensional reconstruction of z-stacks in ImageJ. For cells stained with SMI-32, synapses were analyzed within 15μm from the center of the cell.

For analysis of Hoechst stained cultures, to maximize the likelihood of analyzing inputs innervating a single postsynaptic cell, a 7.5μm radius circle was drawn around the relay neuron nucleus. Only inputs within the circle were analyzed for main figures; axons whose trajectory was tangential to the 15μm circle were discarded. To assess changes in the distribution of boutons from 14 to 18 DIV (see Figures 2c, 3c, S3e), relay neurons were selected for analysis based on proximity to both red and green axons. RGC axons that did not form boutons were included in this analysis. To quantify synaptic competition both via fixed tissue studies and time lapse imaging (Figures 3e, 4), and to determine C1q labeling of competing inputs (Figure 5), only relay neurons that were dually innervated by both a red and a green input were analyzed. Only one red input and one green input were analyzed per relay neuron. If multiple axons crossed the relay neuron, then the input that was selected for analysis was based on the following criteria: 1) boutons that were in closest proximity to the relay neuron in the x-, y- and z-axes, 2) largest number of boutons or biggest bouton(s), and 3) boutons of the competing red and green inputs were in the same focal plane. Number of cells and cultures are indicated for each experimental condition in the figure legends.

### Time lapse imaging

For time lapse imaging, 15-16 DIV cocultures were labeled with Hoechst (1:5000) for 30 min in the incubator. For 12 hour imaging sessions, CNO was added 12 hours prior to imaging as well as into the imaging media for a total of 24 hours of CNO stimulation. For 24 hour imaging sessions, CNO was applied directly to the cultures prior to imaging. Control cultures were unstimulated. Data from 12 and 24 imaging sessions were pooled as no significant difference was observed between the two imaging conditions. Cocultures were placed in an imaging chamber (Tokai-hit) filled with the culture medium which was set at 37°C, 10% CO_2_. Time lapse images were acquired every hour for 12 hours or every 6 hours for 24 hours using an UltraView Vox spinning disk confocal microscope equipped with diode lasers (405nm, 445nm, 488nm, 514nm, 561nm, and 640nm) and Volocity image acquisition software (Perkin Elmer). Z-stacks were analyzed using Image J (NIH). Regions of the dLGN where there were a large number of red and green axons were identified and imaged to maximize the probability of seeing competitive events. Cells of interest were retrospectively identified as those relay neurons that were at some point dually innervated by both a red and a green bouton, as these are the cells that we can observe actively undergoing synaptic competition; those that were only innervated by an input of a single color were discarded. Boutons were scored by eye based on swelling of the axonal segment in close proximity to the relay neuron nucleus. Due to the difficulty of co-labeling for a postsynaptic marker during live imaging, we could not calculate synapse density from the time lapse data. Instead, we quantified the % of relay neurons that received a presynaptic bouton from either a red or green RGC.

### Electrophysiology

Cultures were transferred to a saline solution for recording consisting of (in mM) 125 NaCl, 26 NaHCO_3_, 1.25 NaH_2_PO_4_, 2.5 KCl, 1 MgCl_2_, 2 CaCl_2_, and 25 glucose. Whole-cell voltage clamp recordings of relay neurons were performed using electrodes of 1.2-2.0 MΩ resistance with an internal solution consisting of (in mM) 35 CsF, 100 CsCl, 10 EGTA, 10 HEPES, and 0.1 methoxyverapamil hydrochloride, pH 7.32 using CsOH. For these experiments, the GABA_A_ receptor antagonist bicuculline (20mM, Tocris, MO) was added to the bath solution to block fast inhibitory synaptic currents. Relay neurons were clamped at -70 or +40 mV to record AMPA- or NMDA-receptor mediated currents, respectively, in response to stimulation of the retinal axons with a saline-filled glass pipette, placed toward the periphery of the retinal explant (see Figure 1K). To confirm inward currents at - 70mV were indeed mediated by AMPA receptors, NBQX (5 μΜ, Tocris, MO) was added to the bath solution. To confirm the slow-decaying outward currents at 40mV were through NMDA receptors, CPP (10 μM, Tocris, MO) was added.

To assess DREADD efficacy, RGCs expressing hM3Dq were visually-identified by their green fluorescence and whole-cell current-clamp recordings were made using glass pipettes (6-8 MΩ) with an internal solution containing (in mM): 116 KMeSO4, 6 KCl, 2 NaCl, 20 HEPES, 0.5 EGTA, 4 MgATP, 0.3NaGTP, 10 NaPO4 creatine, pH 7.25 with KOH. After achieving whole-cell configuration, cells were brought to a resting potential between -56 and -65 mV by injecting a constant holding current. 3 sequential square-pulse current injections were then delivered once per minute and adjusted in magnitude so as to elicit action potentials across a range of firing rates. After 10 minutes of consistent baseline firing, CNO (10μM, Sigma-Aldrich, MO) was applied to the bath. Firing rates and resting membrane potentials 8-12 minutes after CNO addition were then compared to baseline values.

### Data representation and statistical analysis

The data were represented as means ± standard error of the mean (s.e.m.). Data were pooled from at least 3 independent experiments. Data were collected and statistically analysed with the researcher blinded to experimental condition. Student’s *t* test was performed for statistical analysis unless otherwise described. Analyses used include one-way ANOVA, unpaired or paired, two-tailed or one-tailed with Tukey’s post-hoc test, Student’s t-test and F test. All p values and samples sizes are indicated in figure legends.

## Acknowledgements

We thank L.V. Goodrich, D.P. Schafer and B. Sabatini for helpful comments on the manuscript and critical discussion. In addition we thank the imaging core at Boston Children’s Hospital including T. Hill for technical support. R.K. was supported by Postdoctoral Fellowship for Research Abroad from Japan Society for the Promotion of Science (JSPS) and B.S. by NIH Grant R01NS07100801A1

## Competing Interests

B.S. is SAB and shareholder in Annexon Biosciences.

---

**Supplemental Figure 1.**
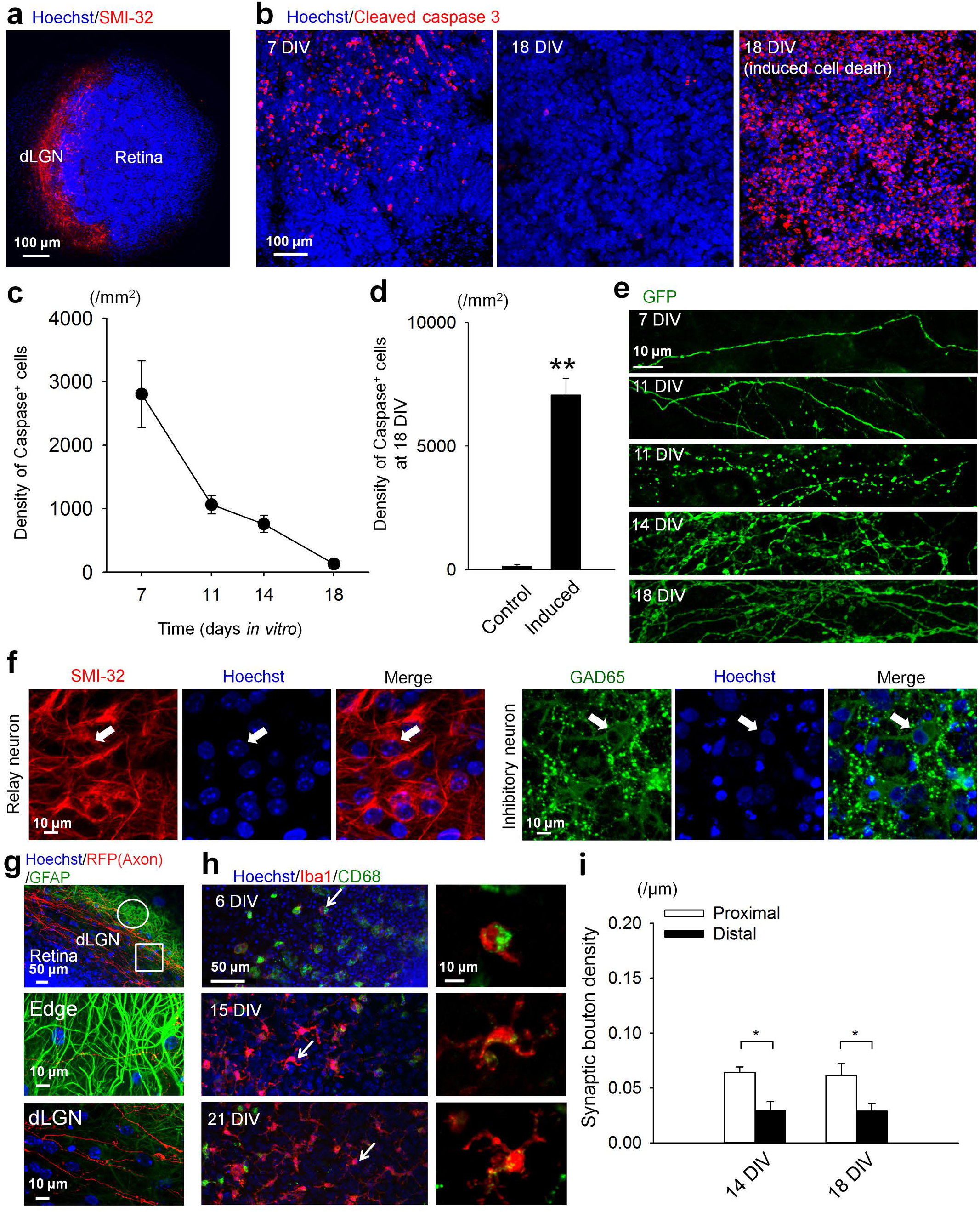
Characterization of the cocultures. (a) Representative coculture stained with the relay neuron marker SMI-32 showing relay neurons in the dLGN region of the culture bordering the retinal explant. (b) Representative images of the retinal region in cocultures immunostained for the cell death marker cleaved caspase-3 at 7, 11, 14, and 18 DIV. As a positive control for cell death, cocultures at 18 DIV were incubated at 4 °C for an hour to induce RGC apoptosis (far right). (c) Time course of the changes in the density of cleaved caspase-3+ RGCs in cocultures at the indicated DIV. (d) At 18 DIV there was very little cleaved caspase-3 staining of RGCs under normal conditions. Induction of apoptosis via incubation at 4°C led to a significant increase in labeling of cleaved caspase-3. Control: n = 4 cultures. Induced: n = 5 cultures. *\*\*P* = 7.89E-06, Student’s *t*-test. (e) Axonal morphologies in the dLGN region of cocultures at 7, 11, 14 and 18 DIV. The two panels at 11 DIV show projecting axons (top) and a blebby, degenerating axon (bottom). No degeneration was observed at 14 or 18 DIV. (f) Representative images of the dLGN *in vitro* at 14 DIV immunostained for the relay neuron marker SMI-32 or the inhibitory neuron marker GAD65. The nuclei were visualized with Hoechst. Arrows indicate relay neuron nuclei (top) or inhibitory neuron nuclei (bottom). (g) Astrocytes are present at the edge of the cocultures. GFAP staining of a coculture showing the retina, dLGN and edge of the culture (left) and magnified images of the dLGN (middle) and edge of the culture (right). Although a few astrocytic processes infiltrate the dLGN, the majority of processes and all the cell bodies are located at the edge of the coculture. (h) Microglia in the cocultures become less phagocytic with time. DIV 6, 15 and 21 cocultures stained for Iba-1 to show microglia cell bodies and processes and CD68, a marker of microglia activation. Note decreased CD68 staining as the cultures mature. Microglia are evenly distributed throughout the dLGN region of the cultures. (i) Few RGC-relay neuron synapses were observed at distal dendrites (>15μm from the soma) compared to proximal dendrites. 14 DIV: n = 29 cells, 11 cultures. 18 DIV: 34 cells, 17 cultures. One-way ANOVA: **P = 0.00120. Tukey-Kramer post hoc test: *\*\*P* = 0.0240 (Proximal vs. Distal, 14DIV), *\*\*P* = 0.0209 (Proximal vs. Distal, 18DIV).

**Supplemental Figure 2.**
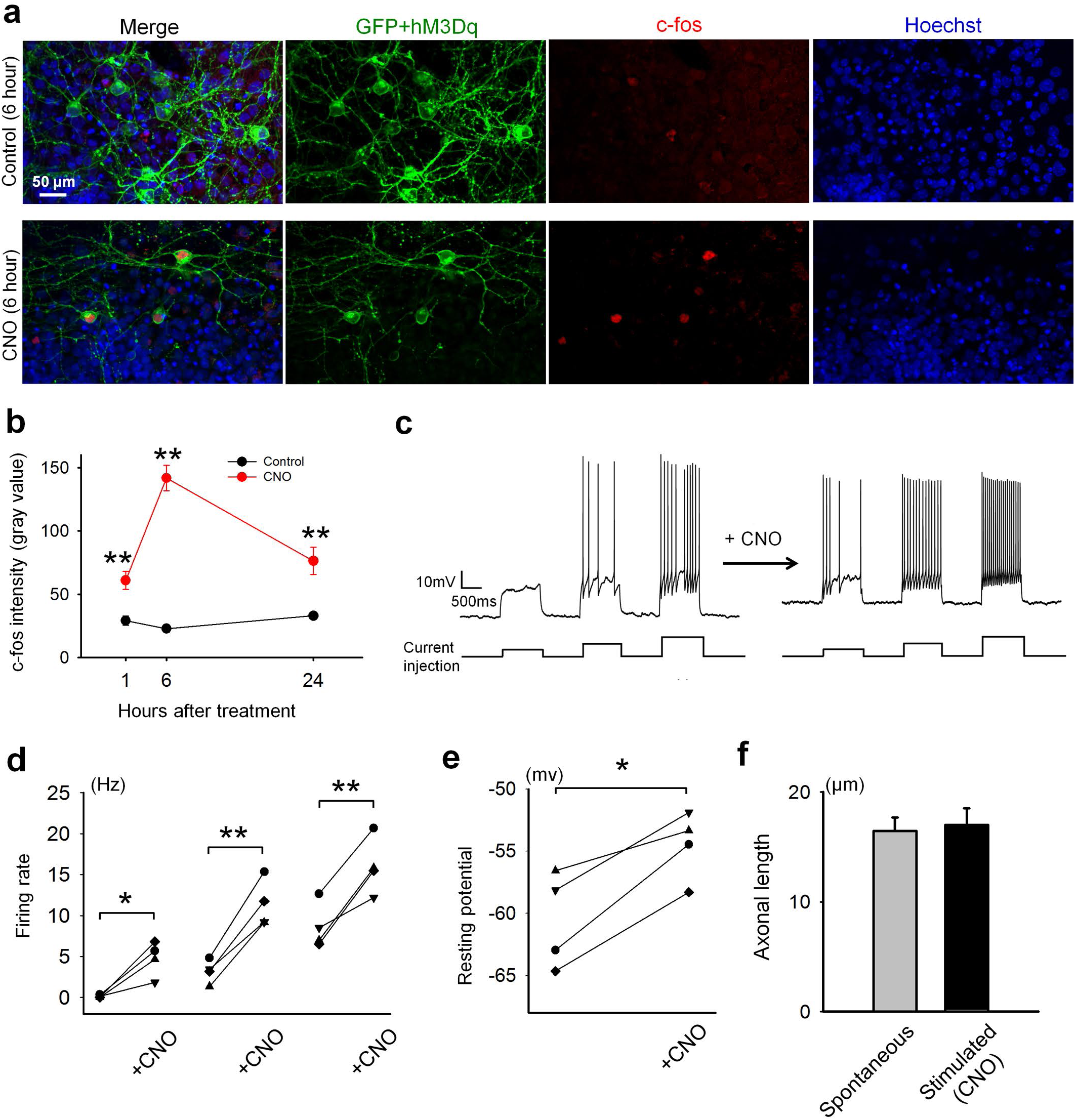
Validation of DREADDs in cocultures. (a) Representative images of control or CNO-treated GFP+/hM3Dq+ RGCs in cocultures immunostained for c-fos at 14 DIV. CNO (10μΜ) was applied for 6 hours before fixation. (b) Time course of c-fos expression (immunofluorescence intensity) at 1, 6, and 24 hours after CNO application at 14 DIV. Per condition: n = 20 cells, 3 cultures. One-way ANOVA: *\*\*P* = 1.28E-23. Tukey-Kramer post hoc test: *\*P* = 0.0222 (1hr control vs. 1hr CNO), **P = 4.93E-14 (6hr control vs. 6hr CNO), *\*\*P* = 0.000438 (24hr control vs. 24hr CNO). (c-e) Electrophysiological recordings from RGCs in cocultures expressing the excitatory DREADD hM3Dq. (c) Example traces before and after bath application of CNO (10uM). Three square-pulse current injections elicit spiking, which is increased upon addition of CNO across all three firing rates. (d) Summary of such recordings from 4 cells. hM3Dq significantly increases firing rates across a range of frequencies. Per stimulation condition: n = 4 cells, 4 cultures. *\*P =* 0.0231 (left). *\*\*P* = 0.00359 (middle), 0.00963 (right). Paired t-test. (e) Mean resting membrane potential before and after CNO application shows a significant depolarization of recorded cells upon hM3Dq activation. n = 4 cells, 4 cultures. *\*P* = 0.0112. Paired *t*-test. (f) CNO application between 14 and 15 DIV did not affect the average axonal length at 18 DIV. Spontaneous: n = 40 axons, 8 cultures. Stimulated: n = 40 axons, 10 cultures. *P* = 0.787, ns. Student’s t-test.

**Supplemental Figure 3.**
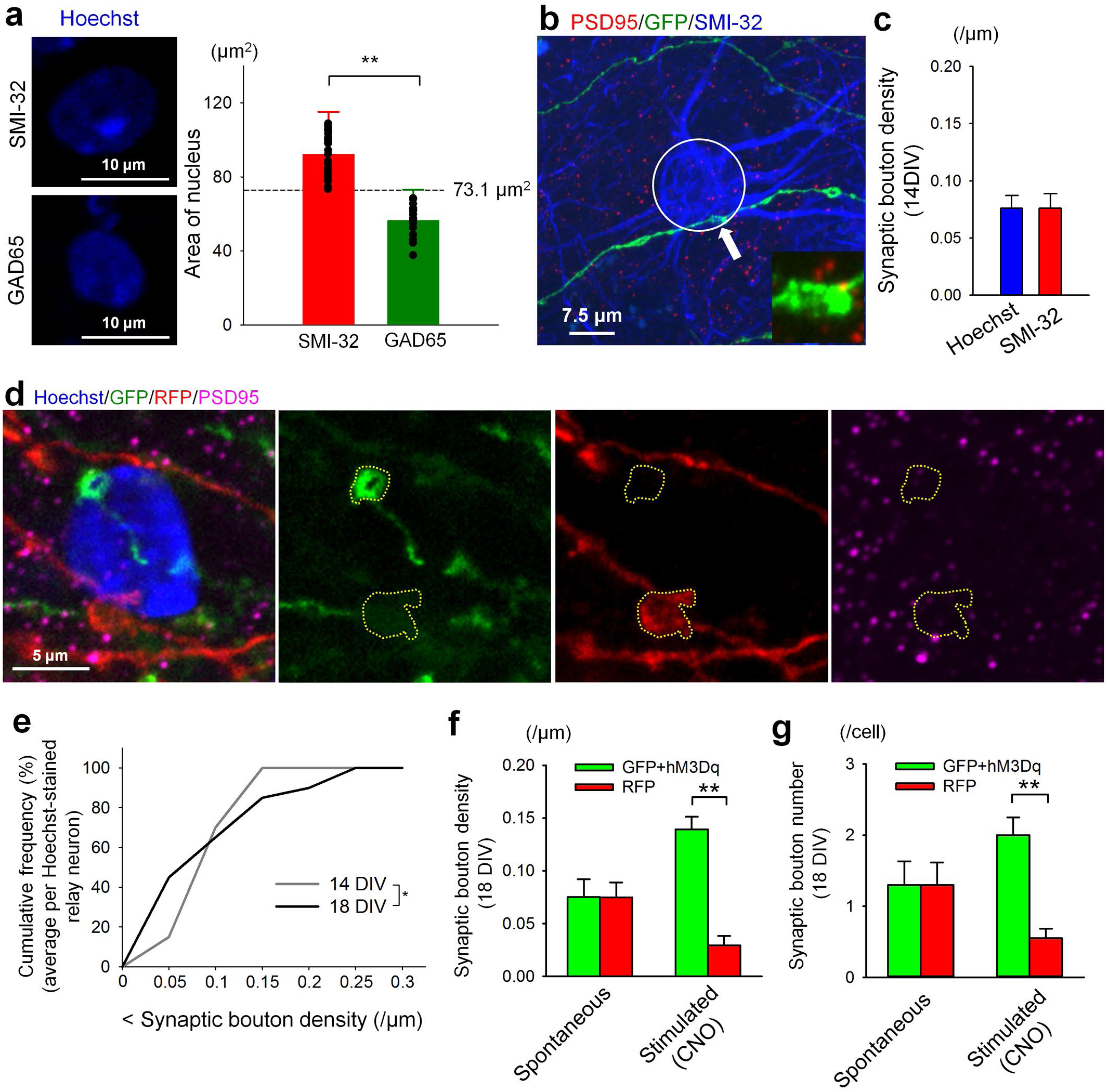
Validation of Hoechst-staining to identify relay neurons. (a) SMI-32+ relay neurons had significantly larger nuclei than GAD65+ inhibitory neurons. Cells with nuclei larger than 73.1μm^2^ (> Mean + 2SD of GAD65 nuclei area) were defined as relay neurons. SMI-32: n = 31 cells, 5 cultures. GAD65: n = 19 cells, 4 cultures. *\*\*P* = 9.043E-16, Student’s *t*-test. (b) As in Figure 1c, except with the 7.5μm radius circle used for analysis of synapses in Hoechst-stained images highlighted. The lower right inset is a magnified image of the PSD95- positive bouton indicated by an arrow. (c) Density of synaptic boutons was analyzed within a 7.5μm radius circle around Hoechst-stained and SMI32-stained neurons. No differences were observed. Per group: n = 21 cells, 8 cultures. *P* = 0.994, Student’s *t*-test. (d) Representative relay neuron contacted by both a GFP+ and an RFP+ input. The relay neuron soma is approximated by the volume of the nucleus, while PSD95+ axonal swellings around the perinuclear region (circled) indicate the presence of synapses in close proximity to relay neuron somas. (e) Cumulative plots of synaptic bouton densities at 14 and 18 DIV from Hoechst-stained neurons. Note the increase in both low and high density inputs at 18 DIV compared to 14 DIV, similar to Figure 2c. 14 DIV: n = 20 cells, 9 cultures. 18 DIV: n = 20 cells, 6 cultures. *P = 0.0112, F-test. (f/g) Under spontaneous conditions, at 18 DIV the density/number of RFP+ and GFP+/hm3Dq+ synapses across all Hoechst-stained cells were not significantly different. Addition of CNO at 14 DIV for 24hrs to activate GFP+/hm3Dq+ neurons led to a significant increase in the density/number of green synapses compared to less active red synapses. Spontaneous: n = 20 cells, 10 cultures. Stimulated: n = 20 cells, 8 cultures. One-way ANOVA: **P = 2.61E-06 in (f). Tukey-Kramer post hoc test: **P = 7.24E-07 (GFP/hM3Dq (CNO) vs. RFP (CNO)) in (f). One-way ANOVA: **P = 0.00415 in (g). Tukey-Kramer post hoc test: **P = 0.00168 (GFP/hM3Dq (CNO) vs. RFP (CNO)) in (f).

**Supplemental Figure 4.**
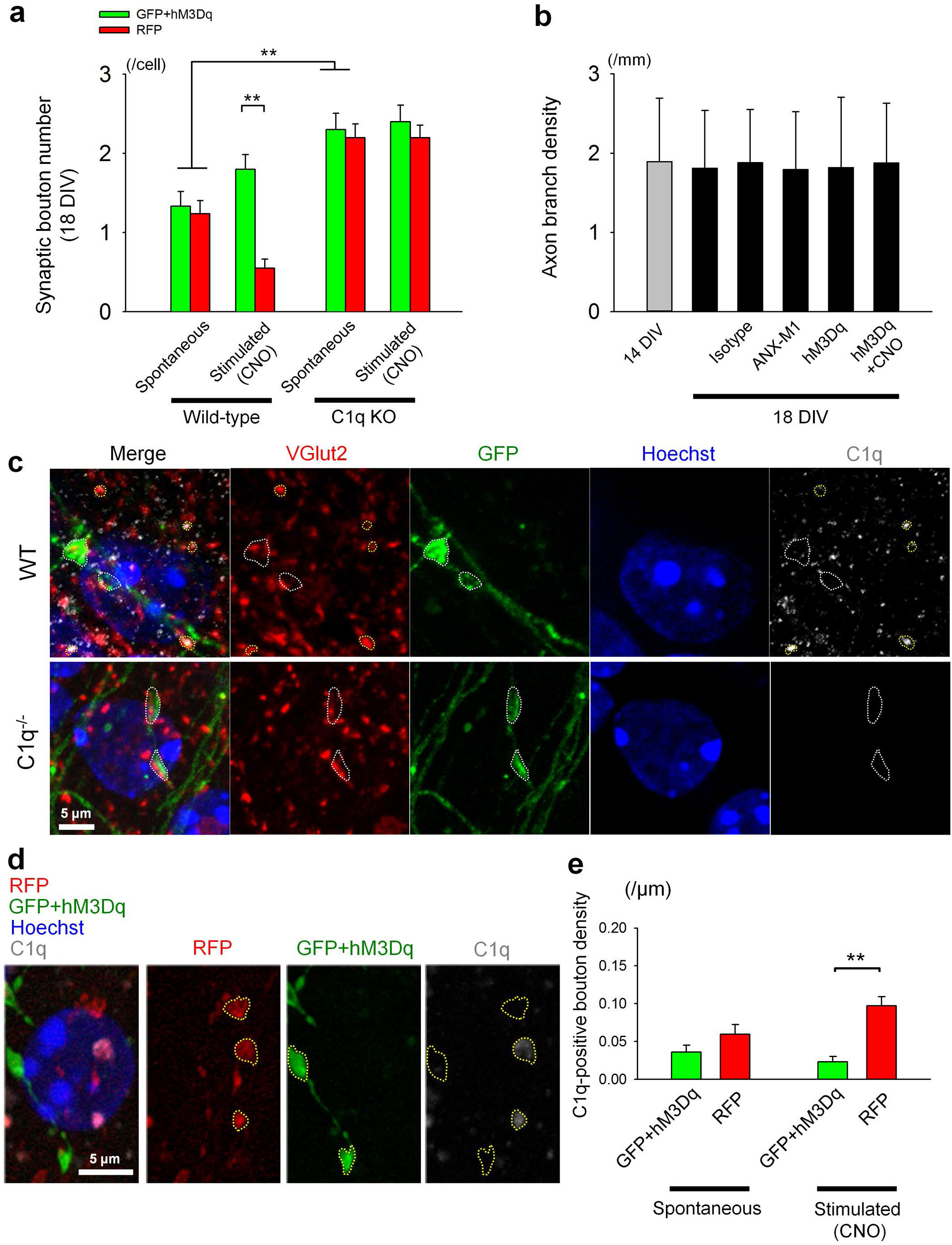
Validation of C1q staining in coculture system. (a) WT and C1q KO cocultures were treated with or without CNO at 14 DIV for 24hrs and fixed at 18 DIV. CNO addition led to a significant increase in the density of green boutons in WT cocultures, but this effect disappeared in KO cocultures. There was also a significant increase in the overall density of red and green boutons formed under spontaneous conditions in KO cocultures compared to WT controls. WT, -CNO: n = 21 cells, 4 cultures. WT, +CNO: n = 20 cells, 4 cultures. KO, -CNO: n = 20 cells, 4 cultures. KO, +CNO: n = 20 cells, 4 cultures. One-way ANOVA: **P = 1.51E-15. Tukey-Kramer post hoc test: **P = 0.00128 (GFP/hM3Dq (WT, -CNO) vs. GFP/hM3Dq (C1q KO, -CNO)), **P = 2.40E-05 (RFP (WT, -CNO) vs. RFP (C1q KO, -CNO)), **P = 2.92E-06 (GFP/hM3Dq (WT, CNO) vs. RFP (WT, CNO)). (b) Axon branch densities were calculated by dividing the total number of axon branches in a field of view by the total axonal length within the field of view. Axon branch density was unaffected by age of cultures or treatment condition. 14 DIV: n = 21 axons, 6 cultures. 18 DIV: n = 22 axons, 7 cultures. Isotype: n = 22 axons, 4 cultures. ANX-M1: n = 21 axons, 4 cultures. hM3Dq: n = 21 cells, 7 cultures. hM3Dq, +CNO: n = 21 axons, 5 cultures. *P* = 0.999, ns, Tukey-Kramer test following One-way ANOVA. (c) The C1qA antibody staining we observed was confirmed to be specific using C1q-/- cocultures. DIV14 WT and C1q-/- cocultures transfected with GFP+/hM3Dq+ were treated with CNO for 24 hours and cocultures were fixed at DIV15. The cocultures were stained for VGLUT2 (Millipore, guinea pig anti-VGLUT2, 1:1000) and C1qA (Epitomics, rabbit anti-C1qA, undiluted culture supernatant). The VGLUT2 staining showed similar intensity between the WT and C1q-/- cocultures; however, in the C1q-/-cocultures, the C1q signal was not detectable using constant settings between WT and C1q-/- cocultures, confirming the specificity of the antibody in this system. In the WT cultures, many C1q puncta could be found colocalized with VGLUT2, consistent with previously reported *in vivo* data (Bialas et al., 2013). (d) Representative confocal images of a Hoechst-stained relay neuron innervated by both GFP+/hM3Dq+ and RFP+ axons at 15 DIV. CNO (10μΜ) was applied at 14 DIV for 18hrs. (e) The density of C1q-positive boutons did not differ between GFP+/hM3Dq+ and RFP+ axons under spontaneous conditions at 15 DIV. C1q preferentially colocalized with non-stimulated RFP boutons at 15 DIV following CNO addition. Spontaneous: n = 26 cells, 5 cultures. Stimulated: n = 26 cells, 5 cultures. One-way ANOVA: **P = 1.71E-05. Tukey-Kramer post hoc test: **P = 2.03E-05 (GFP/hM3Dq (CNO) vs. RFP (CNO)).

**Supplemental Table 1.** Categorization of cells based on innervation status at the start and end of time lapse imaging. After image acquisition, relay neurons near both red and green axons were identified and innervation status was scored at every time point. Innervation status falls into four categories: 1) Both, where both a red and green bouton are present on the relay neuron, 2) Green, where only a green bouton is present, 3) Red, where only a red bouton is present and 4) None, where no boutons are present. Cells that were only innervated by one colored input were not analyzed. The remaining cells that were at some point dually innervated were grouped into 12 categories depending on innervation status at the start and end of the 12- or 24-hour imaging session, e.g., the cells in column 1 began with both a red and green bouton and ended with only a green bouton. To determine whether activity or timing of input arrival biases synaptic competition, cells in the relevant categories (columns) were summed and the percentage of cells was determined per culture. For example, the percentage of cells that began with GFP+ boutons in coculture 1, spontaneous treatment is (2+1+1+2)/26 = 0.23 (sum of columns 5, 8, 9 and 10 divided by total number of cells in coculture 1). These percentages were averaged across cultures within a treatment group (i.e., + or -CNO) and spontaneous and stimulated conditions were compared.

|  | Culture # | 1.<br>Both<br>to<br>Green | 2.<br>Both<br>to<br>Red | 3.<br>Both<br>to<br>Both | 4.<br>Both<br>to<br>None | 5.<br>Green<br>to<br>Green | 6.<br>Red<br>to<br>Green | 7.<br>Red<br>to<br>Red | 8.<br>Green<br>to<br>Red | 9.<br>Green<br>to<br>Both | 10.<br>Green<br>to<br>None | 11.<br>Red<br>to<br>Both | 12.<br>Red<br>to<br>None | Total<br>cells<br>/culture |
| --- | --- | --- | --- | --- | --- | --- | --- | --- | --- | --- | --- | --- | --- | --- |
| Spontaneous<br>Raw #s | 1 | 2 | 3 | 3 | 4 | 2 | 1 | 3 | 1 | 1 | 2 | 2 | 2 | 26 |
|  | 2 | 2 | 1 | 1 | 2 | 2 | 2 | 1 | 0 | 1 | 2 | 4 | 0 | 18 |
|  | 3 | 1 | 2 | 2 | 2 | 4 | 0 | 3 | 1 | 1 | 0 | 1 | 2 | 19 |
|  | 4 | 3 | 1 | 6 | 4 | 0 | 1 | 4 | 0 | 1 | 0 | 2 | 2 | 24 |
|  | 5 | 0 | 6 | 5 | 4 | 1 | 2 | 2 | 0 | 1 | 0 | 1 | 0 | 22 |
| Stimulated<br>Raw #s | 1 | 5 | 0 | 1 | 6 | 6 | 4 | 0 | 1 | 1 | 2 | 2 | 1 | 29 |
|  | 2 | 6 | 3 | 5 | 0 | 1 | 6 | 3 | 1 | 4 | 1 | 0 | 0 | 30 |
|  | 3 | 6 | 2 | 5 |  | 5 | 1 | 2 | 0 | 2 |  | 2 |  | 25 |
|  | 4 | 1 | 1 | 2 | 4 | 7 | 2 | 0 | 0 | 0 | 2 | 1 | 0 | 20 |

**Supplemental Table 1. Categorization of cells based on innervation status at the start and end of time lapse imaging.**
| Figure | 4b: Start |  | 4b: End |  | 4c |  | 4d |  | 4e |  |
| --- | --- | --- | --- | --- | --- | --- | --- | --- | --- | --- |
| Group (x-axis) | GFP | RFP | GFP | RFP | Winner<br>first | Winner<br>second | GFP<br>wins | GFP<br>loses | RFP<br>wins | RFP<br>loses |
| Numerator<br>(column #) | 5, 8, 9, 10 | 6, 7, 11, 12 | 1, 5, 6 | 2, 7, 8 | 5, 7 | 6, 8 | 5 | 8 | 7 | 6 |
| Denominator<br>(column #) | Total cells<br>/culture | Total cells<br>/culture | Total cells<br>/culture | Total cells<br>/culture | 1, 2, 5,<br>6, 7, 8 | 1, 2, 5,<br>6, 7, 8 | 5, 8,<br>9, 10 | 5, 8,<br>9, 10 | 6, 7,<br>11, 12 | 6, 7,<br>11, 12 |

